# Alternating polarity integrates chemical and mechanical cues to drive tissue morphogenesis

**DOI:** 10.1101/2023.08.09.552692

**Authors:** Miriam Osterfield

## Abstract

The spatial patterning of molecules within a sheet of cells directs morphogenesis in many epithelial tissues. In *Scaptodrosophila* follicle cells, Par3/Bazooka and aPKC were previously shown to localize to a set of cell edges destined to elongate and form the base of each eggshell dorsal appendage. This study establishes that the mechanism underlying this localization pattern is an alternating, in-plane polarity of the cells. A candidate screen identified several potential molecular players, whose epistatic relationships were then examined using a custom culture assay. These experiments demonstrated that positive feedback between actin polymerization and PI4P production leads to polarization of these cells individually, while mechanical force coordinates polarization among these cells. This work adds to the growing evidence for a role of mechanics in cell polarity, and also provides an example where morphological differences between species can be understood at the level of changes in fundamental cell biological processes.

**Summary:** In *Scaptodrosophila*, the alternating (left, right, left) polarization in a row of cells drives the formation of up to eight eggshell respiratory filaments. This study uncovers the underlying pathway, which integrates chemical signaling through actin and PI4P with mechanical force.

## Introduction

Epithelial morphogenesis is commonly driven by spatial patterning within the epithelial sheet. This patterning sometimes occurs at the cellular level; for example, apical constriction in a subset of cells can cause dramatic tissue shape changes (Sawyer et al., 2010). However, spatial patterning in other cases is defined subcellularly, at the level of cellular edges. Examples include the orthogonal, planar polarized distributions of myosin and Par3 underlying *Drosophila* germband extension (Bertet et al., 2004, Zallen and Wieschaus, 2004), the curved actomyosin cables which help drive tissue buckling or invagination in multiple systems (Houssin et al., 2020, Laplante and Nilson, 2006, Roper, 2012), and the patterned adhesion molecules which control aspects of cellular architecture in the *Drosophila* eye (Chan et al., 2017, Hayashi and Carthew, 2004).

Subcellularly defined patterns can form in multiple ways. Some patterns are formed on the boundaries between transcriptionally distinct cells; for example, actomyosin cables form on the boundary of Echinoid-expressing and non-expressing cells (Laplante and Nilson, 2006, Laplante and Nilson, 2011, Wei et al., 2005). Other localization patterns can be formed in the absence of sharp expression boundaries, as with the Frizzled/Planar Cell Polarity (PCP) pathway, whose components can localize to edges oriented in a particular direction within a field of essentially identical cells (Butler and Wallingford, 2017). Interestingly, similar spatial patterns can be created by fundamentally different mechanisms. The planar polarized myosin distribution in germband stage *Drosophila* embryo superficially appears quite similar to the PCP pattern of other tissues, but is caused by the interactions between stripes of cells expressing different levels of multiple cell surface proteins (Pare et al., 2019, Pare et al., 2014, Zallen, 2007).

The Drosophilid eggshell, which is molded by the follicular epithelium during late oogenesis, has long been a popular model for studying epithelial patterning and morphogenesis (Berg, 2005, Cetera and Horne-Badovinac, 2015, Horne-Badovinac, 2020, Horne-Badovinac and Bilder, 2005, Montell et al., 2012, Osterfield et al., 2017, Pyrowolakis et al., 2017, Wu et al., 2008). The *Drosophila melanogaster* eggshell has two dorsal appendages (DAs) or respiratory filaments, each formed from a primordium composed of a patch of roof cells bordered to the anterior by a single row of floor cells (Dorman et al., 2004). In the process of DA morphogenesis, the floor cells zipper together into a double row of cells, forming a seam along the bottom of the DA tube (Dorman et al., 2004, Osterfield et al., 2013). In contrast, the eggshell of *Scaptodrosophila lebanonensis* (or its synonym, *Scaptodrosophila pattersoni*, (Bächli et al., 2005)) can have anywhere from 4 to 8 DAs, but all of these structures arise from a single primordium (O’Hanlon et al., 2018, Osterfield et al., 2015). In this case, there is still a single row of floor cells forming the anterior border to a patch of roof cells. However instead of zipping together, the floor cells simply elongate asymmetrically; as a result, the bottom of each DA tube is formed by just two elongated floor cells, with the boundary between them running the entire length of the tube. Prior to morphogenesis, the floor-floor cell edges destined to elongate are marked by an increased localization of Par3 (called Bazooka/Baz in flies) and aPKC. These alternate spatially with floor-floor edges that lack Par3 or aPKC enrichment, and which do not elongate and instead separate neighboring DAs from each other (Osterfield et al., 2015).

In this study, I provide evidence that this alternating pattern of cell edge fate is a consequence of an alternating, planar polarization of the cells, and characterize some key molecular components and mechanical forces driving this phenomenon.

## Results and Discussion

### Alternating edges within a single row of cells show transient F-actin enrichment

As described previously, the edges formed between floor cells in *S. lebanonensis* fall into two classes: edges that are enriched for Par3/Baz and aPKC, spatially alternating with edges that are negative for these markers (Figure 1A-B and (Osterfield et al., 2015)). An antibody-based screen was performed to look for additional molecules that exhibit alternating polarity, initially focusing on candidates that regulate polarity in other systems (Kaplan and Tolwinski, 2010). F-actin (filamentous actin, visualized using phalloidin) was used as a marker for cell boundaries in some of these experiments. Unexpectedly, F-actin showed dramatic enrichment on the Par3-positive edges (Figure 1C).

**Figure 1:**
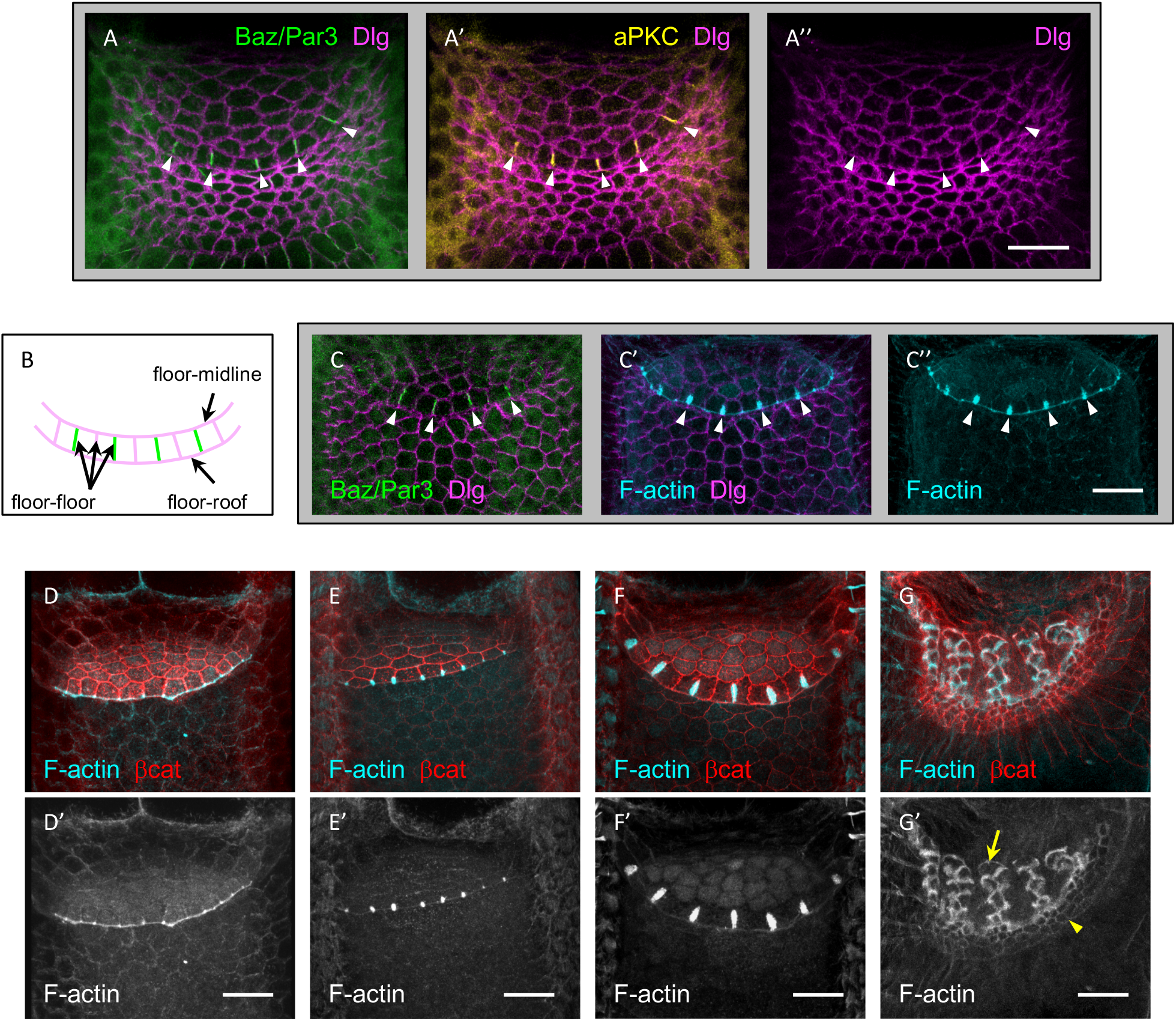
Alternating edges within a single row of cells show transient F-actin enrichment. A-A’’) Baz/Par3 (green) and aPKC (yellow) co-localize to an alternating set of edges (white arrowheads) within the floor cell domain of a stage 10 *S. lebanonensis* egg chamber. Dlg (magenta), a septate junction protein, marks cell outlines. B) Schematic of floor cell domain. The floor-midline domain boundary is located to the anterior (up), and the floor-roof domain boundary to the posterior (down). Alternating floor-floor edges are colored green. C-C’’) F-actin (cyan) is highly enriched on the same alternating set of floor-floor edges (white arrowheads) as Baz/Par3 (green). D-G) Developmentally ordered sequence of stage 10 (D-F) and stage 11 (G) egg chambers, showing dynamics of F-actin localization. β-catenin (red), an adherens junction protein, marks cell outlines. F-actin is initially localized most strongly to the floor-roof boundary (D), then to every floor-floor edge (E), then to alternating floor-floor edges (F). During dorsal appendage formation, F-actin is primarily enriched on roof cell edges (G); levels are higher on roof cells that have already undergone convergent extension and begun forming appendages (yellow arrow) than on those that have not yet (yellow arrowhead). These stages were ordered according to morphological features, including the overall size of the egg chambers (which increases over time) and the shape of the roof cells (which apically constrict during stage 10B, until dorsal appendage formation in stage 11). This ordering was subsequently confirmed by the developmental progression seen with *in vitro* culture assays (see Figure 3). Anterior is up. Scale bars = 20 μm.

The localization of F-actin suggested a possible function in elongating these edges, and/or a function in specifying them. Therefore, I examined the localization of F-actin over a developmental period encompassing both specification and elongation of these edges (Figure 1D-G), corresponding to oogenesis stages 10-12 (Spradling, 1993). The first appearance of a non-uniform membrane distribution for F-actin occurs early in stage 10B, when F-actin is enriched on a smooth, curved boundary that separates the floor cells from the roof cells (Figure 1D). Myosin was also previously shown to accumulate along this border, indicating the presence of an actomyosin cable (Osterfield et al., 2015). At slightly later stages, F-actin becomes enriched on each floor-floor cell edge, beginning along the floor-roof boundary and extending anteriorly (Figure 1E). Still later, but still within stage 10B, this pattern is refined so that only alternating floor-floor edges show prominent F-actin localization (Figure 1F). As the dorsal appendages start to form (stage 11), F-actin continues to localize to alternating floor-floor edges, but also becomes enriched on the membranes of neighboring roof cells (data not shown). As dorsal appendage morphogenesis continues, all of the roof cells edges become enriched for actin, although the roof cells that have intercalated and entered the growing appendages have much higher levels relative to those that have not yet undergone this process (Figure 1G). Strikingly, F-actin is no longer enriched on floor-floor edges at this stage, although these edges continue to elongate (Figure 1G). The later spatial patterns suggest that F-actin may indeed be involved in force generation, for example during early floor-floor edge elongation and/or during apical expansion of the roof cells after incorporation into the dorsal appendage tube. However, the early spatial patterns, in particular the refinement of an every-edge to alternating pattern, suggested that F-actin may also (or alternatively) play a patterning role, by helping specify which set of floor-floor edges will elongate.

### Multiple regulators of the actin cytoskeleton co-localize to the same alternating cell edges

To examine this possibility further, I conducted an immunostaining screen, this time against candidate proteins which interact with or regulate actin. This screen yielded three candidate proteins which co-localize with F-actin to alternating floor-floor edges. Two of these are F-actin-binding proteins which can modulate mechanical or structural properties of the cytoskeleton (Figure 2A-B): α-actinin (Sjoblom et al., 2008) and Sallimus/Kettin, one of two fly homologues of vertebrate Titin (Burkart et al., 2007, van Straaten et al., 1999). The third protein that co-localized with F-actin to alternating floor-floor edges was SCAR (Figure 2C), a WASP-family protein that promotes actin polymerization through the Apr2/3 complex (Pollitt and Insall, 2009).

**Figure 2:**
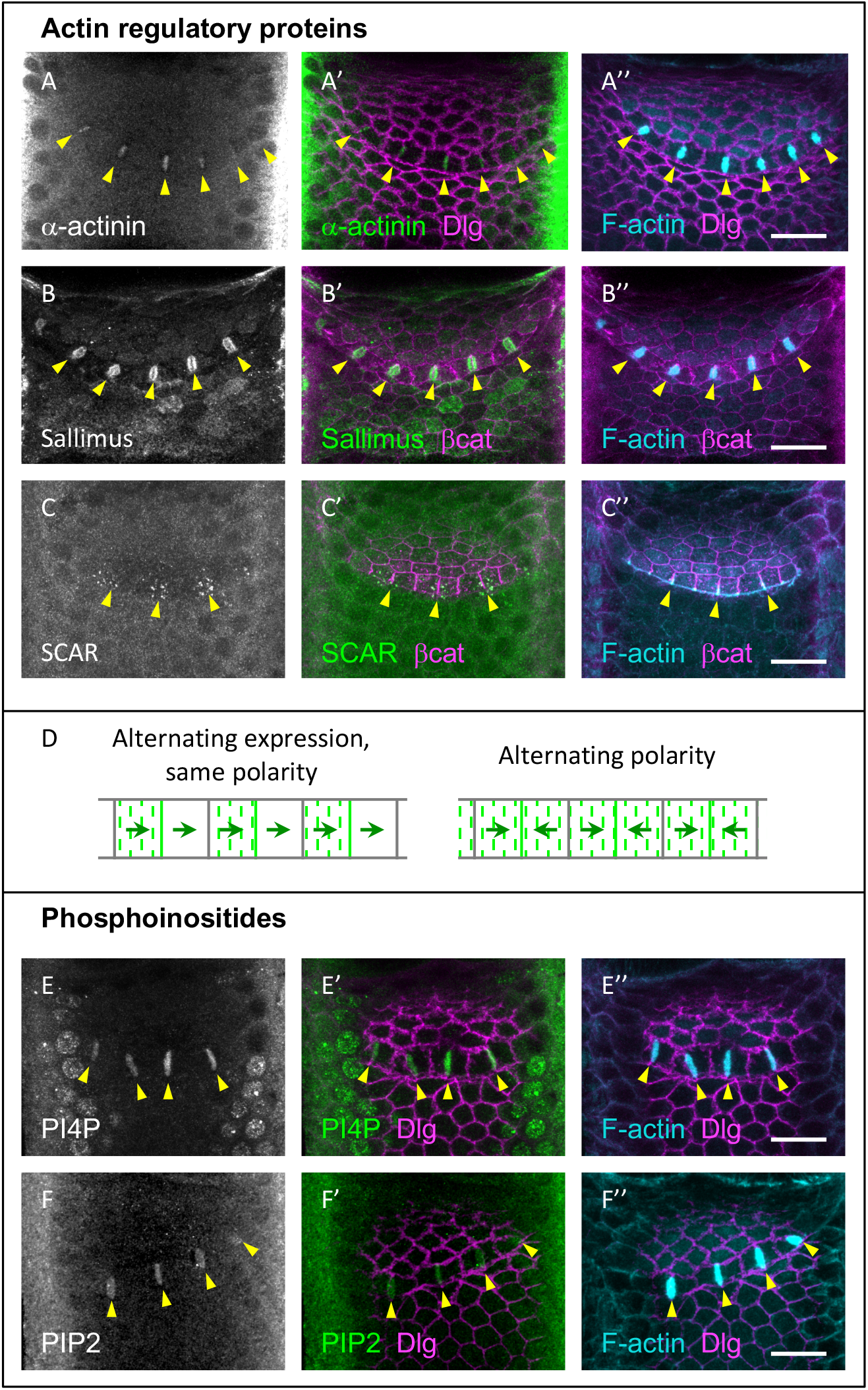
Multiple regulators of the actin cytoskeleton co-localize to the same alternating cell edges. A-C) Immunostaining of candidate actin regulatory proteins α-actinin (A), Sallimus/Kettin/D-Titan (B), and SCAR (C). Dlg or β-catenin (magenta) marks cell outlines. α-actinin and Sallimus co-localize with F-actin (cyan) to alternating edges (yellow arrowheads), while Scar localizes to puncta bordering nascent alternating edges (yellow arrowheads). D) Schematic of two possible underlying causes for the localization of proteins to alternating edges. The localization pattern of SCAR above supports a model invoking alternating polarity (right) rather than alternating expression (left). E-F) Immunostaining of phosphoinositides PI4P (E) and PIP2 (F) reveals co-localization with F-actin (cyan) to alternating edges (yellow arrowheads). Dlg (magenta) marks cell outlines. PIP2 antibody staining may reflect either PI(3,4)P2 or PI(4,5)P2 (see main text). All images in this figure represent at least 3 independent samples with a similar localization pattern. Anterior up. Scale bars = 20 μm.

The localization of SCAR to alternating floor-floor edges is interesting in several ways. Unlike the other candidate molecules described here, SCAR’s alternating pattern is most easily seen in intermediate stages (i.e. between Figures 1E and 1F), when actin is still somewhat enriched on every floor-floor edge, but extends further anteriorly along an alternating subset of edges. This suggests that SCAR may function relatively early in selecting the set of floor-floor edges that will exhibit high levels of actin, aPKC, and the other candidate molecules. Additionally, SCAR-positive puncta cluster on either side of these alternating edges, indicating that SCAR protein in each floor cell is concentrated near one of these edges. This punctate staining in each cell helped resolve a question about the nature of the process specifying these alternating edges. In principle, an alternating set of cell-cell edges could accumulate a specific cell-polarity-regulated protein in two ways (Figure 2D). The cells could all be polarized the same way within the plane of the epithelium, but with only alternate cells expressing the specific protein. Or, the cells could all express the protein, while the polarity of each cell alternates (right, left, right, etc.). The localization pattern of SCAR clearly supports the model of alternating polarity in this system.

SCAR, like other WASP-family members, contains a basic region that can bind phosphoinositides (Pollitt and Insall, 2009). Since phosphoinositides regulate the actin cytoskeleton (Saarikangas et al., 2010) and have been implicated in cell polarity in several other systems (Hammond and Hong, 2018, Mao and Lecuit, 2016), they formed the next group of candidate molecules examined here for localization in *Scaptodrosophila* floor cells. Of the anti-phosphoinositide antibodies used, two showed clear binding to the F-actin-enriched set of floor-floor edges (Figure 2E-F): anti-PI4P, and an anti-PIP2 antibody that binds both PI(3,4)P2 and PI(4,5)P2 (Brzeska et al., 2012), both of which are more highly phosphorylated derivatives of PI4P.

### PI4P and F-actin require each other for localization to alternating cell edges

To test whether and how any of these candidate molecules may function in alternating polarity, it is useful to consider how cell polarity is thought to work in other systems. Cell polarity is generally thought to arise when small initial asymmetries in the localization of key molecules are amplified by positive feedback (Altschuler et al., 2008, Mao and Lecuit, 2016). Although cells can polarize spontaneously, symmetry-breaking signals from the environment often influence the direction of polarization; for example, a gradient of chemoattractant can bias the front-to-rear polarity of a migrating cell, while signaling between neighboring cells biases the direction of polarity on a planar polarized epithelium.

If the alternating polarity seen in *Scaptodrosophila* floor cells works according to similar principles, blocking the localization or activity of key components should result in characteristic phenotypes that reveal the role of the component. For example, removing a positive feedback component should block the polarized accumulation of other molecules, while removing a component that communicates polarity information between cells should randomize the directions in which cells polarize. Therefore, I developed an assay in which one candidate was perturbed pharmacologically, and the resulting localization of other selected candidate molecules was determined. To determine the developmental stage of the egg chambers at the beginning and end of the culture experiment, I took advantage of an unusual feature of *S. lebanonensis* oogenesis: all of the ovarioles in a single female develop synchronously (O’Hanlon, Dev Genes Evol. 2018). Therefore, each experiment was done using a synchronized clutch of stage 10 egg chambers dissected from a single female, divided among three groups. One group was fixed after the dissections were complete, and analyzed later to reveal the pre-culture stage of the egg chambers, while the remaining groups were cultured in experimental (drug) or control (vehicle) containing media before fixation and analysis.

To examine the function of PI4P, this culture assay was performed using a variety of drugs that inhibit either PI kinases or PI phosphatases. One of these drugs, the PI4K/PI3K kinase inhibitor PIK93, produced a striking phenotype. PIK93 inhibits both the development of alternating actin-rich edges when added to samples before the alternating polarity stage (Figure 3A-C), and the maintenance of such edges in samples in which alternating polarity has already been established (Figure 3D-F). The alternating localization of aPKC (Figure 3B’) and PI4P itself (Figure 3E’) were also lost in response to PIK93 treatment in these experiments. These results suggest that PIK93 is acting to prevent the formation of PI4P, and that both F-actin and aPKC require PI4P for their localization to alternating edges.

**Figure 3:**
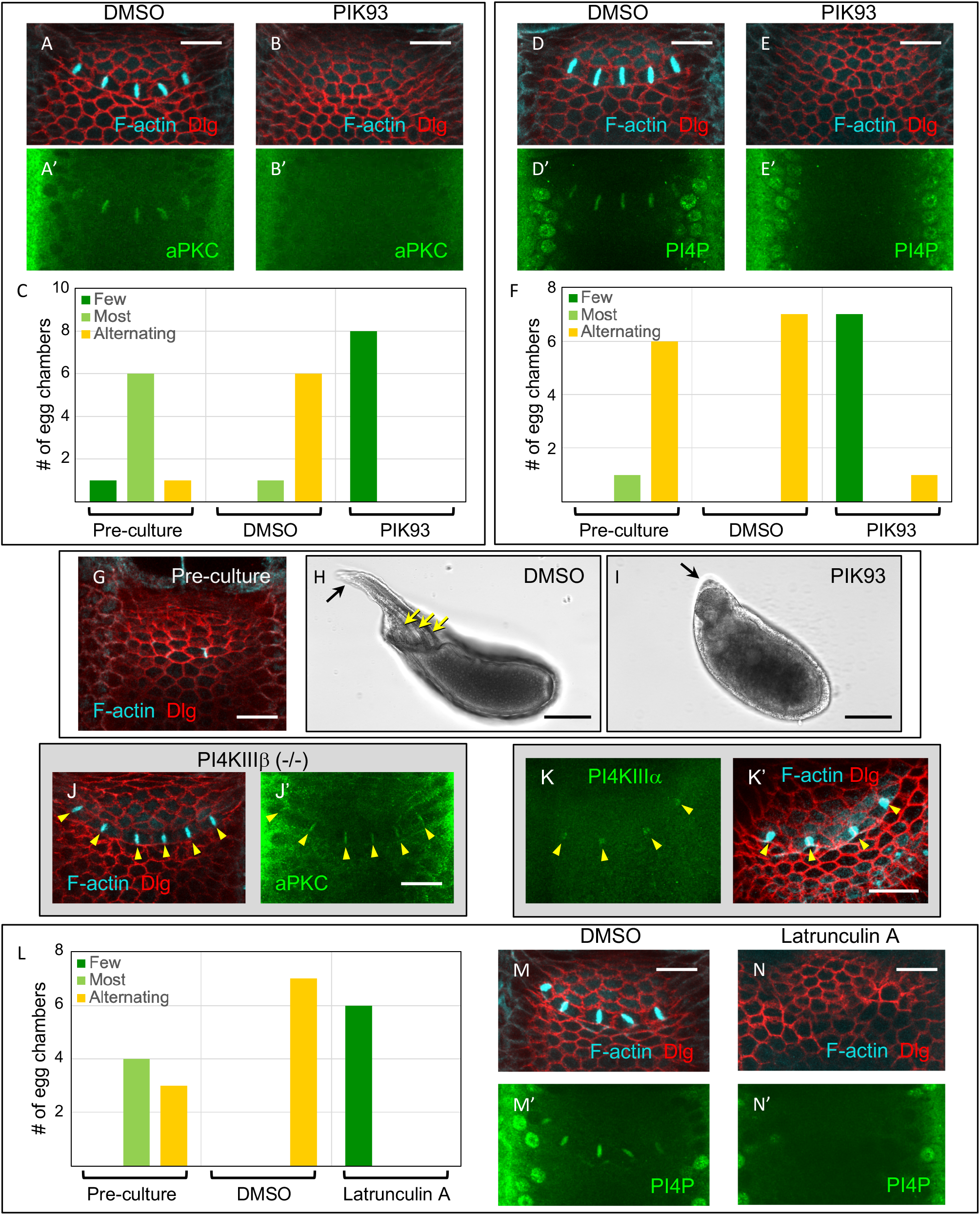
PI4P and F-actin require each other for localization to alternating cell edges. A-C) Culture assay testing the effect of PI4K/PI3K kinase inhibitor PIK93 on the development of alternating polarity. Representative immunostainings (A-B) and quantification (C) are shown. Samples were scored according to F-actin localization pattern; see Supplemental Figure S1 for details. The association between treatment (DMSO versus drug) and outcome (alternating versus non-alternating) is significant (p = 0.0014; two-tailed Fisher’s exact test). D-F) In vitro culture assay testing the effect of PI4K/PI3K kinase inhibitor PIK93 on the maintenance of alternating polarity. PI4P and F-actin mostly fail to localize to alternating edges under PIK93 treatment conditions; the single PIK93-treated sample categorized as “alternating” had both PI4P and F-actin localized to alternating edges. The association between treatment and outcome, calculated as in C), is significant (p = 0.0014). G-I) Representative samples from an overnight culture experiment testing the effect of PIK93 on the development of dorsal appendages. G) Pre-culture samples all had few or no actin-rich edges (n=7/7). H) DMSO treated control samples all had dorsal appendages (n=7/7). Three dorsal appendages (yellow arrows) are visible from this lateral view. I) PIK93 treated samples all lacked dorsal appendages (n=6/6). The association between treatment (DMSO or PIK93) and outcome (dorsal appendages or not) is significant (p = 0.0006; two-tailed Fisher’s exact test). Anterior indicated by black arrow. J) Immunostaining for F-actin and aPKC in *PI4KIIIβ* homozygous mutant egg chambers (yellow arrowheads). Alternating polarity was consistently seen in appropriately staged egg chambers from this genetic background (n=12 egg chambers, from 2 homozygous mutant females). K) Immunostaining for PI4KIIIα protein in wild type *S. lebanonensis* egg chambers. PI4KIIIα co-localizes with F-actin at the alternating polarity stage (yellow arrowheads) (n=10/12). L-N) In vitro culture assay testing the effect of Latrunculin A on alternating polarity. PI4P and F-actin fail to localize to alternating edges under Latrunculin A treatment. The association between treatment and outcome, calculated as in C), is significant (p = 0.0006). Scale bars = 100 μm for H, I. All other scale bars = 20 μm.

To further explore these results, egg chamber culture experiments were also incubated overnight to allow oogenesis to complete (Figure 3G-I). DMSO (solvent control) treated samples developed normally; nurse cells transferred their contents to the oocyte (a process called “nurse cell dumping”), and multiple dorsal appendages formed (Figure 3H). In contrast, PIK93 treated samples failed to form dorsal appendages (Figure 3I), although nurse cell dumping proceeded normally. This result is consistent with the hypothesis that alternating polarity in the floor cells is required for dorsal appendage formation in this species.

PIK93 selectively inhibits the activity of type-III PI4 Kinases, as well as some PI3 Kinases (Balla et al., 2008). There are two type-III PI4 Kinases in D. melanogaster (Balakrishnan et al., 2015), and these both have orthologues in the S. lebanonensis genome. *PI4KIIIβ* (called *four wheel drive*, or *fwd*) is a non-essential gene in *Drosophila* (Brill et al., 2000), suggesting that *PI4KIIIβ* mutant *Scaptodrosophila* might be viable. Indeed, homozygous *PI4KIIIβ* mutant *S. lebanonensis* flies could be generated (see Methods), but protein localization in egg chambers from these mutants appeared entirely normal (Figure 3J), indicating that *PI4KIIIβ* is not required for alternating polarity. Since the other type-III PI4 Kinase, *PI4KIIIα* (or *Pi4KIIIα*) is required for viability in *Drosophila* (Yan et al., 2011), generating homozygous mutants in *Scaptodrosophila* seemed unlikely to work. Instead, I generated an antibody against the *S. lebanonensis* PI4KIIIα protein, and discovered that it co-localizes with F-actin to alternating floor-floor edges (Figure 3K). The combined evidence of its localization pattern and the effect of PIK93 makes PI4KIIIα is an excellent candidate for involvement in alternating polarity. Interestingly, PI4KIIIα has been implicated in cell polarity in several other cell types (Koe et al., 2018, Tan et al., 2014, Yan et al., 2011).

The next candidate examined using the culture assay was F-actin. When added to egg chambers, Latrunculin A disrupts F-actin globally as expected, and in particular, F-actin enrichment on alternating floor-floor edges is lost. Additionally, Latrunculin A also causes the failure of PI4P to localize on alternating edges (Figure 3L-N). Together with earlier results (Figure 3E’), this indicates that PI4P and F-actin each require the other for their localization to alternating floor-floor edges. Crucially, this suggests that PI4P and F-actin are components of a positive feedback loop. However, the manner in which PI4P and F-actin regulate each other’s localization may be indirect. In particular, it is worth noting that PI4P could potentially exert its biological effects by acting as a precursor in the formation of other phosphoinositides.

### Mechanical stress regulates alternating polarity

The identification of a feedback loop involving PI4P and F-actin helps explain how floor cells become polarized, but it sheds no light on why the polarization of these cells alternates. Insight into this question came from an incidental observation: additional actin-rich edges sometimes appeared on ectopic cell edges (floor-midline, midline-midline, or floor-roof) in samples prepared for the culture assay, including in pre-culture and solvent control samples (Figure 4 A-C and data not shown). Furthermore, both PI4P and aPKC localized to these edges (Figure 4B-C), suggesting that they are molecularly equivalent to the actin-rich alternating floor-floor edges seen in tissues prepared for immunostaining only (in Figures 1-2). Unlike egg chambers prepared for simple immunostaining, those prepared for culturing were dissected from the surrounding muscular sheath before fixation. This process can exert mechanical stress on the egg chamber, and the more forceful dissection techniques I initially used when developing this assay seemed to result in more frequent occurrences of ectopic actin-rich edges.

**Figure 4:**
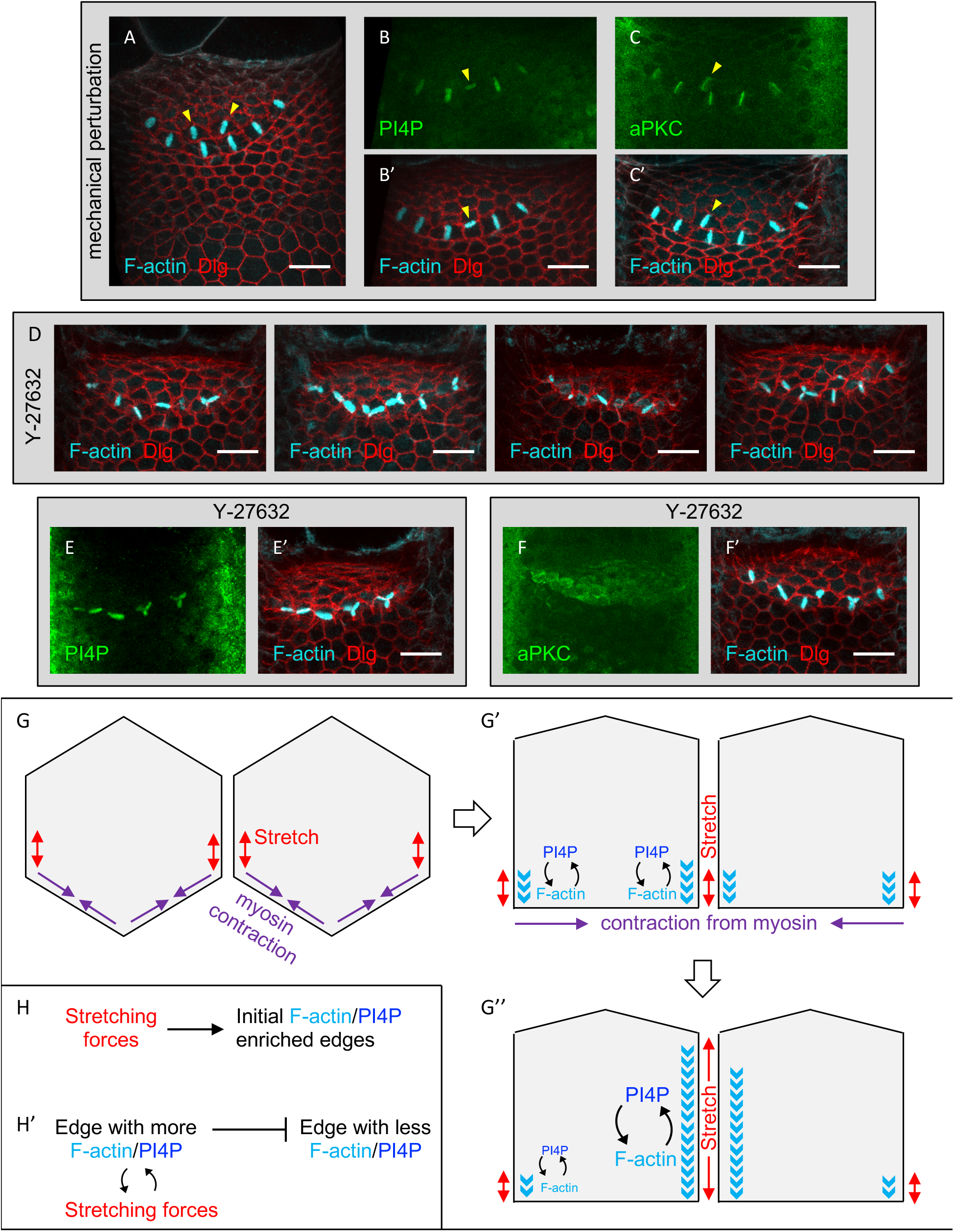
Mechanical stress regulates alternating polarity. A) Control (DMSO) sample from a culture assay, with ectopic actin-rich edges (yellow arrowheads) in the midline cell domain. B-C) Egg chambers that were intentionally subjected to external force by squeezing them through the surrounding muscle during dissection. Ectopic actin-rich edges (yellow arrowheads) are also positive for PI4P (B) and aPKC (C). The highest proportion of egg chambers exhibiting ectopic edges in a single experiment involving intentional squeezing was 4/16; the relatively low penetrance of this phenotype may indicate a requirement for a specific pattern/amplitude of applied force or a specific developmental stage. D) Four representative examples of Y-27632 treated egg chambers from a single experiment. F-actin is enriched on random edges in and near the floor cell domain. E) A representative Y-27632 treated egg chamber, immunostained for PI4P. PI4P co-localizes with F-actin under this treatment. F) A representative Y-27632 treated egg chamber, immunostained for aPKC. aPKC localizes diffusely to the apical surface and fails to co-localize to actin-rich edges. G-H) Model of alternating polarity establishment; see main text. G-G’’ are cartoons of two neighboring floor cells, showing sequential steps of alternating polarity establishment. H is a schematic of the proposed interaction between stretching forces and components of the F-actin/PI4P positive feedback loop during the early phase represented by G-G’. H’ is a schematic of the proposed interactions during the later phase represented by G’-G’’.

The above observations suggested that mechanical force may play some role in polarizing these cells. To investigate further, I tested how the development of alternating polarity was affected by Y-27632, a Rho-Kinase inhibitor that is frequently used in *Drosophila* to block myosin-dependent tension (Vasquez et al., 2016). In egg chamber culture assays, Y-27632 induced a striking phenotype: actin-rich edges were still formed in the floor cell domain, but were found in an apparently randomized distribution (Figure 4D), indicating that polarity is no longer coordinated correctly between neighboring cells. Intriguingly, although PI4P localized to these edges (Figure 4E), aPKC did not (Figure 4F). This implies that aPKC is not required for PI4P and F-actin polarization within individual cells, so it is probably not a component of the PI4P/F-actin positive feedback loop. However, it is unclear whether aPKC localization is required for the communication of polarity information between neighboring cells, or whether its mis-localization in response to Y-27632 treatment is an unrelated effect. Future testing of this point will require the development of either genetic or pharmacological approaches that can acutely disrupt aPKC function in *Scaptodrosophila*.

### Discussion

Together, the observations that increased mechanical stress leads to extra actin- and PI4P-rich edges, and that Y-27632 application with its accompanying decrease in stress leads to randomized actin- and PI4P-rich edges, suggest that mechanical force plays a role regulating the spatial pattern of polarized edges in this system. One model consistent with all of the evidence presented here is that extensive force, i.e. stretching, positively regulates one or more components of the F-actin/PI4P positive feedback loop (Figure 4G-H). In this model, the first event that occurs is the appearance of a myosin cable along the floor-roof boundary (Osterfield et al., 2015). This causes that boundary to constrict, exerting a pulling force on adjoining edges (Figure 4G-G’). This stretching recruits one or more components of the F-actin/PI4P positive feedback loop, causing actin polymerization near the sites of pulling, i.e. the floor-floor edges where they meet the roof domain (Figure 4G’,H). As actin polymerization spreads along each floor-floor boundary, the underlying membrane stretches further, feeding into the F-actin/PI4P/force positive feedback loop. Positive feedback at the fastest growing edge could deplete limiting components elsewhere in the cell, resulting in unidirectional polarity, i.e. a single actin-rich edge per cell (Figure 4G’’,H’); alternatively, an unidentified long-range inhibitory cue might act to suppress F-actin/PI4P accumulation elsewhere in the cell (Altschuler et al., 2008, Mao and Lecuit, 2016). Since faster-growing edges should exert stronger forces on adjacent cells, this mechanism would communicate polarity information between neighboring cells, and thus directly lead to the alternating pattern of cell polarity.

The equivalent alternating polarity pathway coupled to mechanical tension may also exist in other systems where repeating structures are formed from an initially uniform epithelium, such as gut villi or branching structures in the lung. It is interesting to note the final localization pattern might depend on details such as whether the tissue being patterned is essentially 1D (as is the case here) versus 2D or 3D. Examining this question should be an interesting future direction.

## Materials and Methods

### Fly stocks and maintenance

*Scaptodrosophila pattersoni* (National Drosophila Species Stock Center, SKU: 11010-0031.00) were used throughout this study. Since this is recognized as the same species as *S. lebanonensis* (Bächli et al., 2005), the latter name, which is the senior synonym, is used throughout the text. Flies were maintained at room temperature on malt fly food (Archon Scientific #R10101) supplemented with active dry yeast (Red Star or Fleischmann’s). To prevent larvae from escaping, vials were plugged with bonded dense weave cellulose acetate closures, i.e. Flugs (Genesee, cat #49-102).

### Antibodies

Primary antibodies used included the following:

Baz/Par3 (kind gift of Jennifer Zallen (Blankenship et al., 2006), rabbit polyclonal (Figure 1A) or guinea pig polyclonal (figure 1C), 1:250-1:500); aPKC (Santa Cruz Biotechnology, PKCζ antibody (H1): sc-17781, mouse IgG2a, 1:50); Dlg (DSHB (Developmental Studies Hybridoma Bank) # 4F3 anti-discs large supernatant, mouse IgG1, 1:50); β-catenin: (DSHB # N2 7A1 Armadillo supernatant, mouse IgG2a supernatant, 1:20); α-actinin (DSHB # 2G3-3D7 supernatant, mouse IgG2a, 1:10); Kettin/Sallimus/Titin (DSHB # 1B8-3D9 supernatant, mouse IgG1, 1:10); SCAR (DSHB # P1C1-SCAR supernatant, mouse IgG1, 1:50); PI4P (Echelon Biosciences # Z-P004, clone PI4-2, mouse IgM, 8 μg/mL); PIP2 (Abcam # ab2335, clone KT10, mouse IgG2b, 1:100).

The polyclonal anti-PI4KIIIalpha antibody was custom made by ABclonal; it was raised in rabbit against amino acids 1-200 of *S. lebanonensis* phosphatidylinositol 4-kinase alpha (NCBI XM_030526222.1) and affinity purified. Lot# E23940 was used at 1:100 (7.2 μg/ml).

Alexa-Fluor-conjugated secondary antibodies (ThermoFisher) were used at 1:500. For experiments using multiple mouse primary antibodies, isotype specific secondary antibodies (ThermoFisher) were used. Phalloidin (Alexa Fluor 647 conjugated, ThermoFisher # A22287) was used at 6.6 nM or 13.2 nM, and was included along with the secondary antibodies in immunostaining procedures to detect F-actin.

### Immunofluorescence

Immunostaining was done essentially according to standard protocols (Ward and Berg, 2005). Young flies were put in fresh maltose vials with dried yeast and kept at 25°C for 2 to 3 days before dissection. Ovaries were dissected from female flies in Schneider’s Medium (ThermoFisher) supplemented with 17% heat-inactivated FBS (Fetal Bovine Serum, ThermoFisher), and ovarioles were partially separated. Samples were fixed for 20 minutes in 4% paraformaldehyde in PBSTwn (Phosphate Buffered Saline plus 0.1% Tween 20), then washed ∼3x 10 minutes in PBSTwn. Samples were permeabilized by incubation in PBSTwn plus 1% Triton X-100 for 1 hour, and egg chambers were separated by triturating the samples at least 10x through a 1 ml pipette tip near the end of this permeabilization step. Egg chambers were then washed ∼3x 10 minutes in PBSTwn, then incubated in blocking buffer (PBSTwn plus 1% Bovine Serum Albumin) for at least 2 hours at room temperature or overnight at 4°C. Samples were incubated in primary antibodies diluted in blocking buffer (for at least 3 hours at room temperature or overnight at 4°C), then washed ∼3x 10 minutes in PBSTwn. Samples were finally incubated in secondary antibodies diluted in blocking buffer (for at least 3 hours at room temperature or overnight at 4°C), washed ∼3x 10 minutes in PBSTwn, and stored in PBSTwn at 4°C until imaging.

Slight modifications were made to the above immunostaining procedure to improve signal or reduce noise for certain antibodies. For staining with anti-phosphoinositide antibodies, permeabilization and the PBSTwn washes immediately following were done cold (in a 4°C room, and/or using ice-cold buffers). For anti-PI4KIIIalpha staining, the blocking buffer (for blocking, primary, and secondary steps) was supplemented with 5% Normal Goat Serum (Abcam).

### Microscopy and image processing

Fluorescent images were acquired using either a Leica SP8 confocal microscope with a 63x (NA 1.3) glycerol immersion objective or a Zeiss LSM 700 confocal microscope with an EC Plan-Neofluar 40x/1.30 oil DIC M27 objective. All samples were imaged in PBSTwn in a glass-bottom dish (MatTek Life Sciences # P35G-1.5-10-C) Images were adjusted for brightness and contrast in ImageJ/FIJI, and a maximum projection of optical planes showing the apical surface is shown. Due to tissue curvature, the left and right extremes of the images sometimes contain signal from more basal portions of the cells. In some cases (Figures 1A,C,D; 3K; 4A,B), the images were first 3D cropped in Imaris 8.4.2 (Bitplane) to more completely remove signal from overlying tissue (the basal surface, nuclei, etc.) in cases when this would block or overwhelm the apically-localized signal.

Brightfield (DIC) images (Figure 3H-I) were acquired on a Zeiss LSM 700 confocal microscope using a Plan-Apochromat 20x/0.8 M27 dry objective and ZEN Blue software.

### Culture assay

To enrich for stage 10B egg chambers, young flies (less than 2 or 3 days post-eclosion) were transferred to fresh malt vials sprinkled with dried yeast and kept at 25°C for about 2.5 days. Female flies were then dissected in Schneider’s Medium (ThermoFisher) supplemented with 17% heat-inactivated FBS (ThermoFisher). A pair of ovaries from a single female was then chosen based on apparent stage (ideally early-to-mid 10B) for a culture assay. Individual egg chambers were separated and removed from the muscle sheath, and any egg chambers that appeared damaged or noticeably different in stage from the rest were discarded. Once all the individual egg chambers to use were selected, they were divided into three roughly equal groups: one to fix immediately, one to culture with the drug being tested (diluted in Schneider’s Medium + 17% FBS), and one to culture with the appropriate solvent control (diluted in Schneider’s Medium + 17% FBS).

Samples to be cultured were placed in 150 μl culture media within the 10 mm well of a glass bottom dish (MatTek). The dishes were covered, and the samples were examined approximately every 15-20 minutes to check developmental progress. All samples were simultaneously removed from culture and fixed (see immunostaining procedure) as soon as one or more egg chambers (in either drug or control condition) appeared to have begun nurse cell dumping (stage 11). After fixation, samples were washed 4x in PBSTwn, then stored in PBSTwn at 4°C overnight, or up to several days, before proceeding with the rest of the immunostaining protocol.

For PIK93 (Sigma SML0546), stock made was 25 mM in DMSO, and final concentration used was 250 μM. For Latrunculin A (Sigma # L5163), stock was 10 mM in DMSO, and final concentration was 50 μM. For Y27632 (Millipore Sigma # 688000), stock was 100 mM in water, and final concentration was 0.2 mM.

### CRISPR injection protocol

To collect embryos for injection, flies were put in collection cages, mounted on collection plates that contained banana media (Archon Scientific #W210) with a smear of yeast paste. Flies were kept on a 12 hour light / 12 hour dark cycle at 25°C, and embryo collections were done within about 5 hours after the beginning of the dark cycle. Zero-to-two hour old embryos were collected, dechorionated in bleach for about 3 minutes, washed thoroughly in water, then mounted on a slide using tape glue (Kiehart et al., 2007) and covered with Halocarbon oil 700 (Sigma #H8898). Injection needles were pulled from quartz glass capillaries with filaments (Sutter Instrument Company # QF100-70-10) using a Sutter Model P-2000 pipette puller with the following program: Heat = 800, Filament = 4, Velocity = 55, Delay = 128, Pull = 110. Needles were loaded using microloader tips (Eppendorf # 5242 956.003), opened by breaking at the tip under Halocarbon oil 700, and used in conjunction with a FemptoJet 5247 Microinjector (Eppendorf) to inject posterior ends of the prepared embryos.

### CRISPR genome modification design

In *D. melanogaster, PI4KIIIβ* (*fwd*) has two alternative start sites, with the start for the fwd-PC isoform falling within the second coding exon for the other isoforms (fwd-PA, PB, PD, and PE.) Publicly available genomic sequence for S. lebanonenesis includes a predicted cDNA sequence (XM_030530475.1) homologous to the longer isoforms of *PI4KIIIβ* (*fwd*), and a genomic scaffold (NW_022060742.1) to which this cDNA can be mapped. Examination of these sequences reveals a similar exon-intron structure as seen in D. melanogaster. In particular, an alternative start codon in the second coding exon (corresponding to the start codon for the fwd-PC isoform in D. melanogaster) appears to be conserved.

To create a frameshift mutation expected to completely disrupt the *S. lebanonenesis PI4KIIIβ* (*fwd*) gene, guide RNAs were designed to target the second coding codon, after this potential alternative start. This location is well upstream of the kinase domain. The CRISPOR tool (Concordet and Haeussler, 2018) was used to choose the following guide + *PAM* as a target: TT GGT CAC TTG GGA AAC TGG *AGG*. The gRNA was made by in vitro transcription using a Hiscribe T7 high yield kit (New England Biolabs #E2040s) using the following primers (Integrated DNA Technologies): Forward primer Sleb_fwd_g5_f: 5’-GAA ATT AAT ACG ACT CAC TAT AGG TTG GTC ACT TGG GAA ACT GGG TTT TAG AGC TAG AAA TAG C-3’; Reverse primer T7_CRISPR_b: 5’-AAA AGC ACC GAC TCG GTG CCA CTT TTT CAA GTT GAT AAC GGA CTA GCC TTA TTT TAA CTT GCT ATT TCT AGC TCT AAA AC-3’. The gRNA was purified using a Microelute RNA clean UP kit (Omega Biotek #6247-01). The following mixture was then prepared for injection into embryos: 2 μg/μl Cas9-NLS protein (Integrated DNA Technologies #1081058), 150 mM KCl, 0.4 μg/μl gRNA. Green food dye (Kroger) was added at approximately 1:10 to help with visualization during embryo injection. Preparation of this injection mixture was based on previously published protocols (Nyberg et al., 2020, Stern et al., 2017).

Injected flies and their progeny were screened for deletions in the second coding exon by conducting PCR on genomic DNA using the following pair of primers: Forward primer fwd_exon2_3f: 5’-GAC AAT GAG CAA TTT TGC TTC ACG-3’ Reverse primer fwd_exon2_3b: 5’-CAA TGA GTT GGC TCT AGC ACA CGC-3’ Heterozygotes to be used in crosses were identified by using the T7 endonuclease I assay (Lin et al., 2014, Nyberg et al., 2020). To genotype individual flies used for immunostaining, PCR was done on genomic DNA from the carcasses and analyzed by Sanger sequencing. The PI4KIIIβ homozygous mutants analyzed here had an 8 bp deletion expected to cause a frame shift in both predicted isoforms of this gene. The following is the sequence of the second coding exon of PI4KIIIβ, with the 8 bp deleted region indicated in capital letters: 5’-a tcc atg ggc ata ttg cta cct cca GTT TCC CAa gtg acc aat aca cgc atc aat cac aca caa cat cat cgg aat cgc agc ctc gac agt gcc ctg cag cgc ata cca gag -3’

## Data availability

The images that were quantified for Figure 3 are available from the corresponding author upon reasonable request. The data underlying Figures 1, 2, and 4 are available in the published article.

## Acknowledgements

I am grateful to Konstantin Doubrovinski, Celeste Berg, Michael Buszczak, and members of their labs for discussion, to Shelby Blythe, David Stern, Tamas Balla, and Jen Liou for technical advice, and to Trudi Schüpbach for comments on the manuscript. This work was funded by NSF grant 1755015.

---

**Figure S1:**
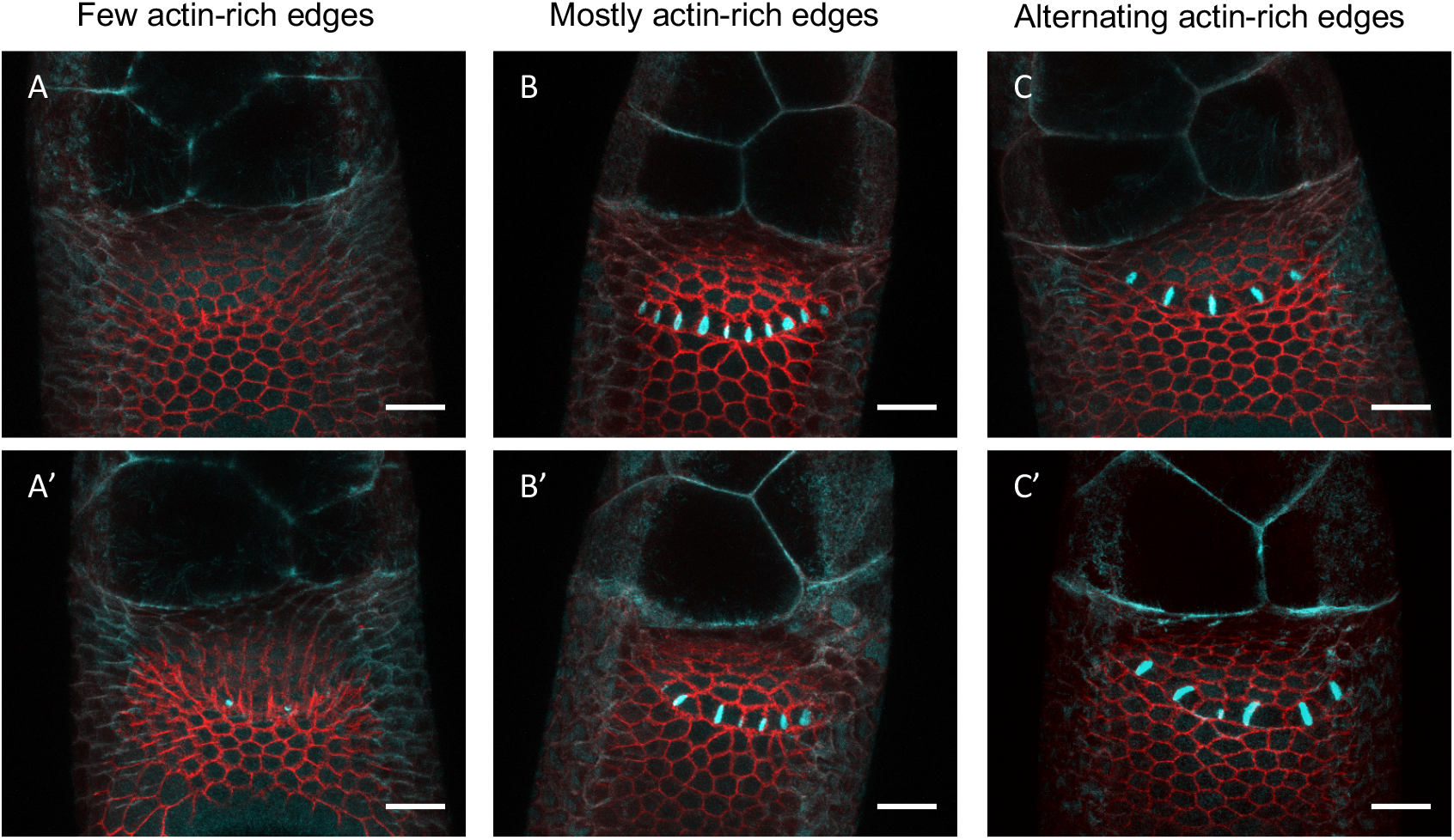
Scoring of culture assay samples. A-C) Examples of samples from culture assays to illustrate scoring scheme. Culture samples were scored according to F-actin localization and placed in one of three categories; those with few or no actin-rich floor-floor edge (Figure S1A-A’, compare to Figure 1 D), those in which most or all floor-floor edges were actin-rich (Figure S1B-B’, compare to Figure 1 E), and those in which alternating floor-floor edges were actin-rich (Figure S1C-C’, compare to Figure 1 F). Note that while panels A, B, and C have actin localization patterns that exactly match the stages presented in Figures 1D, 1E, and 1F (with no, all, or alternating actin-rich edges respectively), panels A’, B’, and C’ can be easily classified as members of the same categories, with one or two “defects”.

